# CCDC86/Cyclon is a novel Ki-67 interacting protein important for cell division

**DOI:** 10.1101/2022.07.01.498427

**Authors:** Konstantinos Stamatiou, Aldona Chmielewska, Shinya Ohta, William C Earnshaw, Paola Vagnarelli

## Abstract

The chromosome periphery is a network of proteins and RNAs that coats the outer surface of mitotic chromosomes. Despite the identification of new components, the functions of this complex compartment are poorly characterised. In this study we identified a novel chromosome periphery-associated protein CCDC86/cyclon. Using a combination of RNAi (RNA interference), microscopy and biochemistry, we studied the functions of CCDC86/cyclon in mitosis. CCDC86/cyclon depletion resulted in partial disorganisation of the chromosome periphery with partial alterations in the localization of Ki-67 and Nucleolin and the formation of abnormal cytoplasmic aggregates. Furthermore, CCDC86/cyclon-depleted cells displayed errors in chromosome alignment, altered spindle length and increased apoptosis. These results suggest that, within the chromosome periphery, different subcomplexes that include CCDC86/cyclon, Nucleolin and B23 are required for mitotic spindle regulation and correct kinetochore-microtubule attachments, thus contributing to chromosome segregation in mitosis. Moreover, we have identified CCDC86/cyclon as a MYC-N regulated gene whose expression levels represent a powerful marker for prognostic outcomes in neuroblastoma.

**Summary statement:** Here we report the identification of CCDC86/cyclon as novel component of the perichromosomal layer. CCDC86 is important for chromosome segregation and represents a strong prognostic marker for neuroblastoma patients.

## Introduction

As cells enter mitosis, chromatin undergoes a remarkable series of structural changes that lead to chromosome condensation and the formation of individual mitotic chromosomes (Paulson et al., 2021) (Booth and Earnshaw, 2017). The composition and function of specific mitotic chromosome domains such as the chromosome scaffold, centromere/kinetochore and telomeres have been extensively investigated. However, the ill-defined compartment known as the chromosome periphery (CP) or perichromosomal sheath/layer is less well characterized. The perichromosomal layer coats the outer surfaces of individual mitotic chromosomes (Dimario, 2004) (Hernandez-Verdun and Gautier, 1994) (Van Hooser et al., 2005) and constitutes approximately one third of the protein mass of mitotic chromosomes (Booth et al., 2016) (Booth and Earnshaw, 2017). This chromosome compartment appears at prometaphase and disappears at telophase when the nuclear envelope reforms (Montgomery, 1900).

Four different functions have been proposed for the chromosome periphery: a) involvement in the maintenance of mitotic chromosome structure, b) establishment of a physical barrier protecting mitotic chromosomes from cytoplasmic proteins following nuclear envelope breakdown, c) protecting chromosomes from sticking to one another and d) acting as a scaffold to distribute nucleolar proteins required for post-mitotic nucleolar reactivation (Montgomery, 1900) (Metz, 1934). Even though experimental evidence is available to support each of these hypotheses, the functional significance of the chromosome periphery remains elusive.

Most chromosome periphery proteins are associated with the nucleolus or the nucleoplasm during interphase until the transition from G_2_ to mitosis when, upon nuclear envelope breakdown, they accumulate at the chromosome periphery in a sequential manner. Among the CP components there are many ribonucleoproteins (RNPs), RNAs (Gautier et al., 1992) (Hernandez-Verdun and Gautier, 1994) (Van Hooser et al., 2005) and the nucleolar proteins fibrillarin, nucleolin, B23/nucleophosmin, peripherin and Ki-67 (Booth and Earnshaw, 2017) (Hernandez-Verdun and Gautier, 1994) (Booth and Earnshaw, 2017; Jansen et al., 1991) (Tollervey and Kiss, 1997) (Yasuda and Maul, 1990).

Despite the fact that the chromosome periphery received little attention other than in studies of nucleolar inheritance (Gautier et al., 1992; Hernandez-Verdun, 2006; Hernandez-Verdun and Gautier, 1994; Hernandez-Verdun et al., 2013; Muro et al., 2010) in the past several years interest in this structure has substantially increased. Ki-67 is one of the earliest proteins associated with the chromosome periphery from early prometaphase to telophase (Van Hooser et al., 2005) and is essential for formation of the chromosome periphery (Booth et al., 2014); in fact, Ki-67 depletion led to the disappearance of the chromosome periphery with significant effects on the distribution of nucleolar components at mitosis (Booth et al., 2014). Interestingly, mitotic chromosomes lacking the perichromosomal layer tend to aggregate, suggesting the interesting hypothesis that Ki-67 acts as a steric and electrostatic coating on chromosomes by forming a sort of biological surfactant important to maintain the individuality of each mitotic chromosome (Cuylen et al., 2016). Although Ki-67 seems to be a master organizer of this compartment, many other proteins are present in the layer. Whether they fulfill specific functions in chromosome structure or dynamics is not generally known, although the recent discovery of the NWC complex and demonstration that it is required for Aurora B recruitment to chromosomes (Fujimura et al., 2020) raises the possibility that other components may also have important roles in chromosome segregation.

Several studies have analyzed the effects of nucleolin and B23/nucleophosmin depletion in human cells. These led to diverse phenotypes, including defects in chromosome alignment and segregation with micronucleus formation, mitotic delay, increased apoptosis and abnormal nucleolar architecture (Ugrinova et al., 2007) (Amin et al., 2007) (Ma et al., 2007). In contrast, fibrillarin depletion did not induce the same phenotype but seemed to be involved in maintaining nuclear shape integrity (Amin et al., 2007).

Late in mitosis, the nucleolar proteins associated with the chromosome periphery become diffuse in the cytoplasm and associate in cytoplasmic particles called nucleolus-derived foci (Hernandez-Verdun, 2011; Hernandez-Verdun et al., 2013) (Muro et al., 2010) (Dundr et al., 2000). Afterwards components of the NDFs fuse to form pre-nucleolar bodies (PNBs), which are distributed throughout the early G1 nuclei (Dimario, 2004; Dundr et al., 2000; Olson and Dundr, 2005). These dynamics are important for the reformation of a functional nucleolus.

The diverse mitotic phenotypes associated with perturbations of the perichromosomal layer raise the possibility that sub-compartments of this structure may have specific mitotic functions. It is therefore important to be able to identify and dissect the various sub-complexes to understand how these chromosome structures are coordinated.

In this study we have identified CCDC86/cyclon as a new component of the perichromosomal layer. We have shown that its depletion causes defects in chromosome alignment and segregation without perturbing the localization of Ki-67 at the chromosome periphery in early mitosis and alters the composition of NDF in anaphase/telophase. This, together with the discovery of the NWC complex (Fujimura et al., 2020) reveals that indeed some subcomplexes of this layer may have specific mitotic functions. We have also discovered that CCDC86 is a MYC regulated whose expression levels are of prognostic value in Neuroblastoma patients.

## Materials and Methods

### Cell Culture, Cloning, and Transfections

HeLa Kyoto cells were grown in DMEM supplemented with 10% foetal bovine serum (FBS) and 1% Penicillin–Streptomycin (Invitrogen Gibco) at 37□°C with 5% CO2.

CCDC86 was cloned in pEGFP N1 and C1 by PCR from HeLa cell cDNA using the following primers:

Fw CGGATCCGGCGGGATGGATACACCGTTAAGG and Rev CAGAATTCGGATCTTGGCTGC

The oligo resistant mutation was created as follows: AGATTCTCCCAGATGTTACAAGAC

For the siRNA treatments, HeLa cells in exponential growth were seeded in 6-well plates, transfected using Polyplus JetPrime (PEQLAB, Southampton, UK) with the appropriate siRNA oligonucleotides (50 nM), and analysed at the times indicated in the experiments. The siRNAs were obtained from Sigma: Control: 5’-CGUACGCGGAAUACUUCGA-3’; CCDC86 Oligo 1 SASI_Hs01_00120887 and CCDC86 Oligo 2: SASI_Hs01_00120888. For the rescue experiments, 400 ng of the wt or oligo-resistant constructs were used.

TET21-N cell line (Lutz et al., 1996) was kindly supplied by Prof Arturo Sala (Brunel University London). Neuroblastoma cells were cultured in DMEM supplemented with 10% FBS and 100 units ml^−1^Penicillin/Streptomycin. TET21-N cells were routinely maintained in medium containing 0.2 mg ml^−1^ G-418 (11811031, GIBCO) and 0.15 mg ml^−1^Hygromycin B (10453982, Invitrogen), and to switch off MYCN expression cells were cultured in the presence of 1 µg ml^−1^ doxycycline (D9891, Sigma-Aldrich).

### Pull down experiments and immunoblotting

GFP or GFP:CCDC86 (400 ng/each) were transfected in HeLa cells. 7×10^6^ cells were collected and snap-frozen in liquid nitrogen. Cell were then resuspended in 200 ml of lysis buffer (25mM TRIS-HCl pH7.4, 150 mM NaCl, 0.1% SDS, 0.5% NP40, 0.5 mM EDTA), incubated on ice for 30’, sonicated and the lysate was cleared by high speed centrifugation.

The supernatant was then incubated with 30 ml of GFP-binder beads (Chromoteck) for 1h at 44^°^C. The beads were washed 4 times in lysis buffer, re-suspended in 2X Laemmli (Laemmli, 1970), boiled and load on a 10% SDS PAGE gel.

For immunoblotting, whole cell lysates were loaded onto polyacrylamide gels. SDS-PAGE and immunoblotting was performed following standard procedures.

### FACS analysis

Control or CCDC86 depleted HeLa cells were subjected to cell-cycle analysis by FACS. Briefly, 1 × 10^6^ cells were fixed with cold ethanol (70%) for 1 hr, centrifuged, and resuspended in PBS containing RNase A (0.2 mg/ml) and Propidium Iodide (10 μg/ml) (Fisher Scientific, P3566). Following a 20-min incubation, cells were analysed using FACS.

The FL2 channel was used to analyse 20,000 events per condition. Gated cells were manually categorized into cell-cycle stages. Cells were analysed following knock-down periods of 48 and 56 hr.

### Antibodies

The primary antibodies were used as follows: Ki-67 (mouse monoclonal BD Transduction laboratory, Oxford, UK) 1:100; nucleolin (rabbit polyclonal; Abcam) 1:300; anti-alpha-tubulin antibody (B512; SIGMA, Gillingham, UK), anti-GFP (Invitrogen, Paisley, UK); Lamin A+Lamin C (ab108595, Abcam) 1:2500.

### Indirect Immunofluorescence

For immunofluorescence, cells were fixed in 4% PFA and processed as previously described (Vagnarelli et al., 2011). Fluorescence-labelled secondary antibodies were applied at 1:200 (Jackson ImmunoResearch). 3D data sets were acquired using either 1) a cooled CCD camera (CH350; Photometrics) on a wide-field microscope (DeltaVision Spectris; Applied Precision) with a NA 1.4 Plan Apochromat lens. The data sets were deconvolved with softWoRx (Applied Precision). 3D data sets were converted to Quick Projections in softWoRx.; or 2) a wide-field microscope (NIKON Ti-E super research Live Cell imaging system) with a numerical aperture NA 1.45 Plan Apochromat lens. The data sets were deconvolved with NIS Elements AR analysis software (NIKON). Three-dimensional data sets were converted to Maximum Projection using the NIS software. In both cases the images were exported as TIFF files, imported into Adobe Photoshop and imported into Inkscape for final presentation.

Live cell imaging was performed with a DeltaVision microscope as previously described (Vagnarelli et al., 2011).

### In silico Neuroblastoma analyses

The analyses of Neuroblastoma data sets were conducted using the R2: Genomics Analysis and Visualisation Platform (http://r2.amc.nl)

### qPCR

Total RNA of TET21-N cells (Control and 24 h doxycycline) was extracted using the Monarch Total RNA Miniprep Kit (New England Biolabs, Hitchin, UK), from which complementary DNA (cDNA) was synthesised by reverse transcription using RevertAid RT Reverse Transcription Kit (Thermo Fisher Scientific, Waltham, MA USA). qPCR was performed using Maxima SYBR Green/ROX qPCR Master Mix (2X) (Thermo Fisher Scientific, Waltham, MA USA) by the Thermo Fischer Scientific QuantStudio 7 Flex Real-Time PCR Instrument. Relative gene expression was calculated using the comparative Ct method (ΔΔCt). GAPDH was used as endogenous reference gene.

qPCR was conducted using the following primers

GAPDH:

Fw: ACCACAGTCCATGCCATCAC

Rev: TCCACCACCCTGTTGCTGTA MYCN:

Fw: CACAAGGCCCTCAGTACCTC

Rev: ACCACGTCGATTTCTTCCTC

CCDC86/Cyclon:

Fw: TTCCTCTCCTGTCGTTCCTT

Rev: AGCGAAAAGGTTCTTCATCC

### In silico Neuroblastoma analyses

The analyses of Neuroblastoma data sets were conducted using the R2: Genomics Analysis and Visualisation Platform (http://r2.amc.nl)

### Statistical analyses

Statistical analyses were performed using a Chi-square test or Fisher exact test or student’s T test.

## Results

### CCDC86/Cyclon is a novel perichromosomal layer component

In order to understand the composition and the function of the perichromosomal layer in mitosis, we need to be able to identify each subcomplex and assess the specific contribution. We have previously shown that depletion of Ki-67 causes the most dramatic phenotype as it is at the top of this hierarchy by controlling the assembly of the entire chromosome periphery (Booth et al., 2014). To identify novel proteins that could be part of this structure and classify them into different subcomplexes, we reanalysed the mitotic chromosome proteome (Ohta et al., 2010; Ohta et al., 2011). As shown before, *bona fide* mitotic chromosome proteins cluster when the proteome is analysed using the “ Abundance” on chromosomes vs “ Enrichment “ classifiers (Ohta et al., 2011). Ribosomal and nucleolar proteins have a lower retention coefficient, possibly indicating significant binding to chromosomes from cytosol during the incubation. This could reflect the nature of these complexes, which undergo major dynamic re-localisation transitions during mitosis.

Based on these two classifiers, we identified as a novel “ bona fide” chromosomal protein the Coiled-Coil Domain Containing 86 (CCDC86/cyclon) (Figure 1 A, B). The gene ontology for this protein includes 3 cellular components: nucleus (GO:0005634), nucleoplasm (GO:0005654) and nucleolus (GO:0005730). Using the Data Visualization for Protein-Protein Interactions (GPS-Prot http://gpsprot.org/), we analysed the known interactome for CCDC86. CCDC86 shows interactions with known nucleolar proteins that belong to the mitotic perichromosomal layer, including Ki-67, fibrillarin, NOP56, and also with histones (Figure 1 C). We therefore generated GFP-fusion constructs (both N and C terminus tag) of CCDC86 for localisation studies.

**Figure 1.**
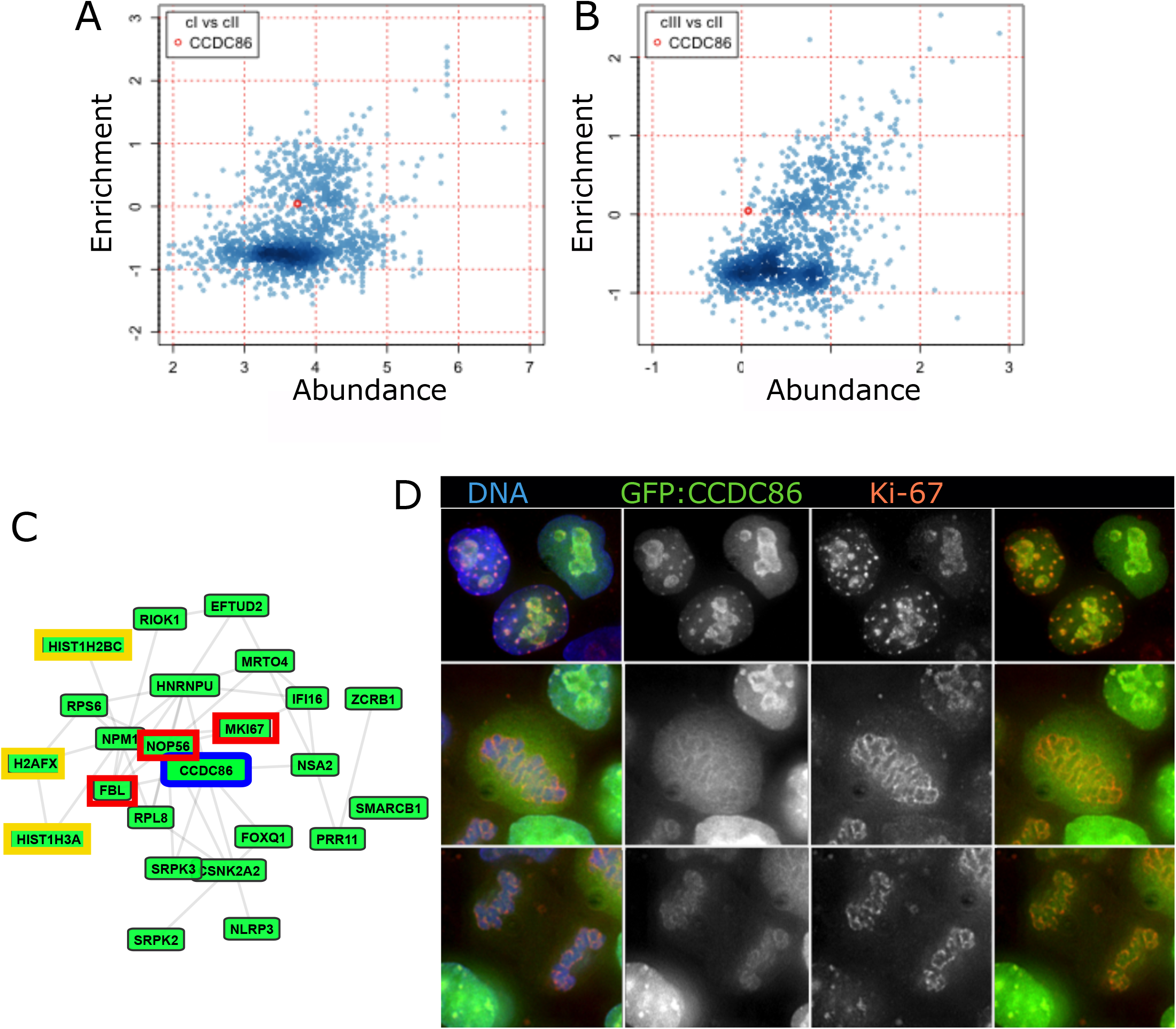
CCDC86/cyclon is a nucleolar protein associated with the chromosome periphery. A, B) 2D scatter graph plotting the fold-change (log2) of protein abundance on mitotic chromosomes (cI, A) or in vitro exchange on chromosomes (cIII, B) versus enrichment on chromosomes (cII) generated in previous proteomics analysis (Ohta et al., 2010). The red circle indicates CCDC86. The color intensities, ruled by the smooth density plot in R, indicate the population densities. C) CCDC86 interactome (GPS-Prot – Human PPIs). In red are highlighted known nucleolar proteins and in yellow histone proteins. D) A GFP:CCDC86 construct (green) was transfected in HeLa cells. Cells were fixed and stained for Ki-67 (red). Representative images showing CCDC86 localisation in interphase (top row), metaphase (middle row) and telophase (bottom row). Blue (DNA).

In HeLa cells, both fusion proteins localised to the nucleolus in interphase. They were mainly dispersed in the cytoplasm in mitosis but enriched at the periphery of the chromosomes. They were most highly enriched at the chromosome periphery during anaphase (Figure 1 D). Interestingly, co-staining with Ki-67 antibodies revealed that the two proteins always co-localise in interphase and mitotic exit but only partially in prometaphase/metaphase (Figure 1 D middle panel). Therefore, this novel protein could be an interesting candidate to dissect individual functions of the chromosomal periphery subcomplexes.

### CCDC86/Cyclon enrichment to the chromosomes depends on Ki-67 and its first AT-hook like domain

As most perichromosomal layer proteins depend on Ki-67 for their localisation (Booth et al., 2014), we tested whether Ki-67 RNAi would also impair the localisation of CCDC86. As predicted, Ki-67 RNAi strongly diminished the enrichment of CCDC86 at the periphery of the chromosomes both in early mitosis and during mitotic exit (Figure 2 A). Therefore, CCDC86 recruitment to the chromosomes is mediated or stabilised by Ki-67. However, the interactome analysed (Figure 1C) showed that histones are also part of CCDC86 network. Moreover, the domain analyses of CCDC86 revealed the presence of 3 conserved AT-hook like motives in the protein (Figure 2B). These motifs are found in a variety of DNA binding and DNA remodelling proteins including the high mobility group (HMG) proteins and the hBRG1 (SWI/SNF) remodelling complex protein that have a preference for A/T rich regions(Reeves and Nissen, 1990). We therefore tested if these motifs were involved in the localisation of CCDC86.

**Figure 2.**
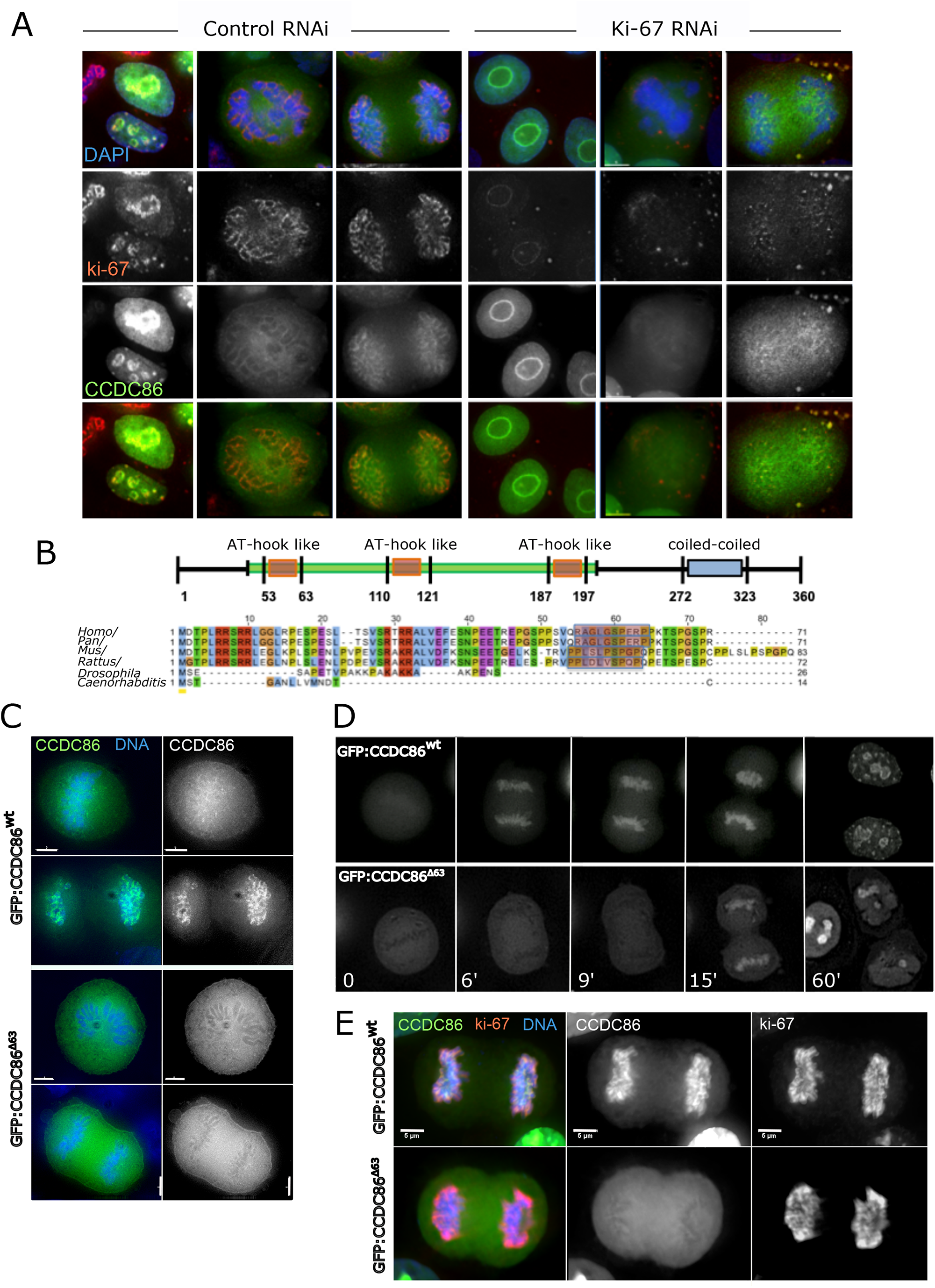
CCDC86/Cyclon requires Ki-67 and the first AT-hook domain for its recruitment to the mitotic chromosomes. A) HeLa cells were transfected with a GFP:CCDC86 construct (green) and treated with control or Ki-67 siRNA oligos. Cells were then fixed and stained for Ki-67 (red) and DNA (Blue). Representative images of interphase (left panels), prometaphase (middle panel) and anaphase (right panel) cells are shown for both Control RNAi and Ki-67 RNAi. B) Top: Scheme of the CCDC86 protein showing the position of the 3 AT-hook like domains and the coiled-coiled region; Bottom: alignment of the N-terminal region of the protein. Shadowed in red are the sequences of the first AT-hook. C) HeLa cells were transfected with either a GFP:CCDC86wt construct of the mutant lacking the first 63 amino acids (GFP:CCDC86D63) (green) then fixed 24 h post transfection. A representative metaphase and anaphase are shown. Scale bars 5 μm. D) Live cell imaging of HeLa cells transfected with either a GFP:CCDC86wt construct of the mutant lacking the first 63 amino acids (GFP:CCDC86D63). E) HeLa cells were transfected with either a GFP:CCDC86wt construct of the mutant lacking the first 63 amino acids (GFP:CCDC86D63) (green) then fixed 24 h post transfection and stained for Ki-67 (red). A representative metaphase and anaphase are shown. Scale bars 5 μm.

We generated GFP tagged truncated proteins by deleting either the first, the first and second and all three AT-hook like domains of CCDC86. All these mutant fusions abolished the localisation of the protein to the chromosomes when analysed as transient transfections in HeLa cells (Figure 2C). In order to gain more information about the dynamic behaviour of the proteins, we conducted live cell imaging using either GFP-tagged CCDC86 wild type protein (GFP:CCDC86^wt^) or a mutant version lacking the first AT-Hook (GFP:CCDC86^Δ^_63_). While GFP:CCDC86^wt^ protein was first seen on the chromosomes in metaphase cells and then strongly accumulated on anaphase chromosomes (Figure 2 D-top panels), the mutant counterpart failed to localise on the chromosomes in metaphase/anaphase (Figure 2 D-top panels). GFP:CCDC86^Δ^_63_ accumulated on chromosomes only at 15 minutes into mitotic exit. This is the time when the nuclear envelope is reforming, and nuclear import starts. Later in G_1_ (60’), the mutant protein accumulated in the nucleoli. This data suggests that, although Ki-67 is necessary for CCDC86 recruitment to the chromosome periphery, it is not solely sufficient as the first AT-hook like domain of CCDC86 is needed for the chromosomal localisation even if Ki-67 is present (Figure 2 E). Alternatively, the N terminal region of CCDC86 contains the domain necessary for the interaction with either Ki-67 or other proteins that are Ki-67 dependant for their recruitment to the chromosomes.

### CCDC86/Cyclon depletion causes the formation of abnormal Nucleolar derived foci (NDF)

We next analysed the mitotic phenotypes of cells in which CCDC86 was depleted. We used two siRNA oligos that can efficiently deplete the protein at 48h (Figure 3A and B) and first analysed the localisation of both Ki-67 and Nucleolin. Interestingly, both proteins were present at the mitotic chromosome periphery both in early mitosis and mitotic exit, but they were also observed in cytoplasmic foci that resemble NDFs in both stages of mitosis. Although Nucleolin is found in NDF foci in normal cells (Dundr et al., 2000), the foci were much bigger and more abundant after CCDC86 depletion and were also present in prometaphase/metaphase (Figure 3 D), which is not normal. Remarkably, after CCDC86 depletion, KI-67 was also detected in NDFs. This is never observed in unperturbed cells. We therefore quantified the frequency of telophase cells in which Ki-67 was present in the abnormal NDFs using lamin A/C co-staining in order to score cells all at the same stage of mitotic exit. The results using two independent siRNAi oligos revealed a significant increase of telophase cells with NDFs containing Ki-67 after CCDC86 depletion (Figure 3E and F). We also wanted to investigate if the reported interactions between Ki-67 and CCDC86 could be confirmed by immunoprecipitation. We used asynchronous HeLa cells transfected with GFP:CCDC86 and pulled down the protein using GFP:trap. By immunoblot analyses we could confirm that GFP:CCDC86, but not GFP alone, was able to pull-down Ki-67 (Figure 3G), thus confirming the interaction *in vivo* between the two proteins.

**Figure 3.**
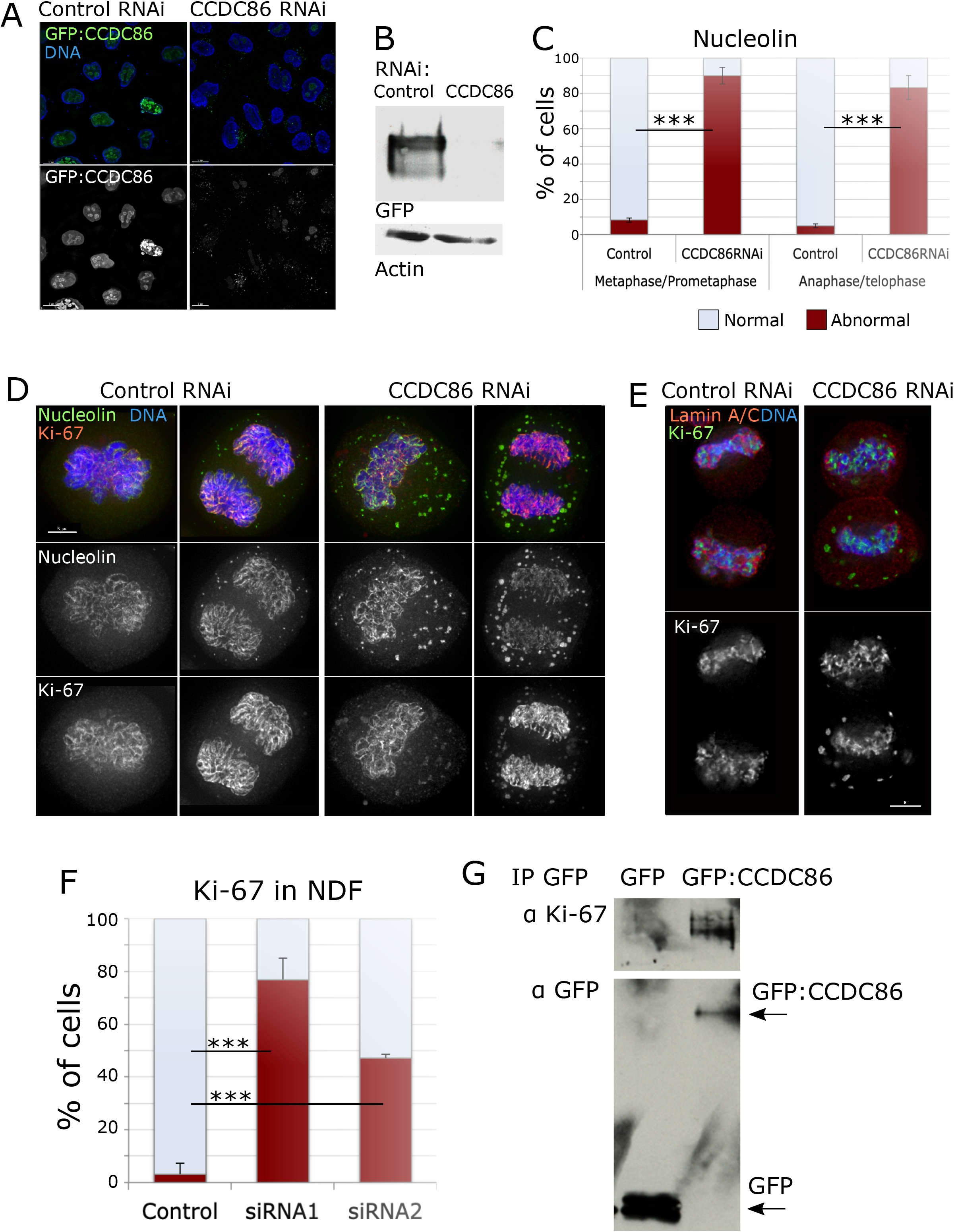
Depletion of CCDC86/Cyclon compromises the mitotic localisation of nucleolin and Ki-67. A) HeLa cells were transfected with a GFP:CCDC86 construct (green) and treated with control or CDCD86 RNAi oligo (#1) for 48h. Cells were fixed and counterstained with DAPI (blue); Representative images showing cells with GFP:CCDC86 (green) in control RNAi and no GFP signal in CCDC86 RNAi. B) SDS PAGE of HeLa cells treated as in (A) and immunoblotted using either GFP (top panel) or beta actin (bottom panel) antibodies. C) Quantification of cells displaying an abnormal nucleolin pattern in metaphase/prometaphase or anaphase/telophase from the experiment in (D). The graph shows the average or 3 independent experiments and the error bars represents the SD. 50 cells from each replica and condition were analysed. Statistical analyses were conducted using the Chi squared test and *** indicates p < 0.0001. D) HeLa cells were treated with either siRNA control or CCDC86 oligos for 48h and the fixed and stained for Nucleolin (green), Ki-67 (red) and counterstained with DAPI (blue). Representative images of a metaphase cell (left) and anaphase cell (right) after each RNAi treatment. Scale bar 5μm. E) HeLa cells were treated with either siRNA control or CCDC86 oligos (Oligo #1 or oligo #2) for 48h and then fixed and stained for Ki-67 (green), LaminA/C (red) and counterstained with DAPI (blue). Representative images of telophase cells (as judged by the recruitment of Lamin A/C around the chromosomes) after each RNAi treatment. Scale bar 5μm. F) Quantification of cells displaying an abnormal Ki-67 pattern in telophase from the experiment in (E). The graph shows the average or 3 experiments and the error bars represent the SD. Statistical analyses were conducted using the X^2^ test and *** indicates p < 0.0001. G) HeLa cells were transfected with either GFP or GFP:CCDC86 constructs. After 24h cells were lysed and pull downs with GFP-trap was conducted. The beads were then boiled, subjected to SDS PAGE and probed with antibodies against Ki-67 or GFP.

### CCDC86/Cyclon is important for chromosome segregation

Having identified CCDC86 as a novel component of the chromosome periphery whose depletion does not abolish the localisation of known perichromosomal layer proteins such as Nucleolin and Ki-67, we wanted to check if depletion of this protein was associated with chromosome defects in mitosis. We first analysed the Fluorescence activated cell sorting (FACS) profile of cells depleted of CCDC86 using the two different oligos at 48 and 56 h. The analyses showed an increase in the sub-G_1_ population (indication of apoptosis) and a decrease in the G_1_ and G_2_ populations with both oligos (Figure 4A). This could suggest that cells lacking CCDC86 enter apoptosis after cell division.

**Figure 4.**
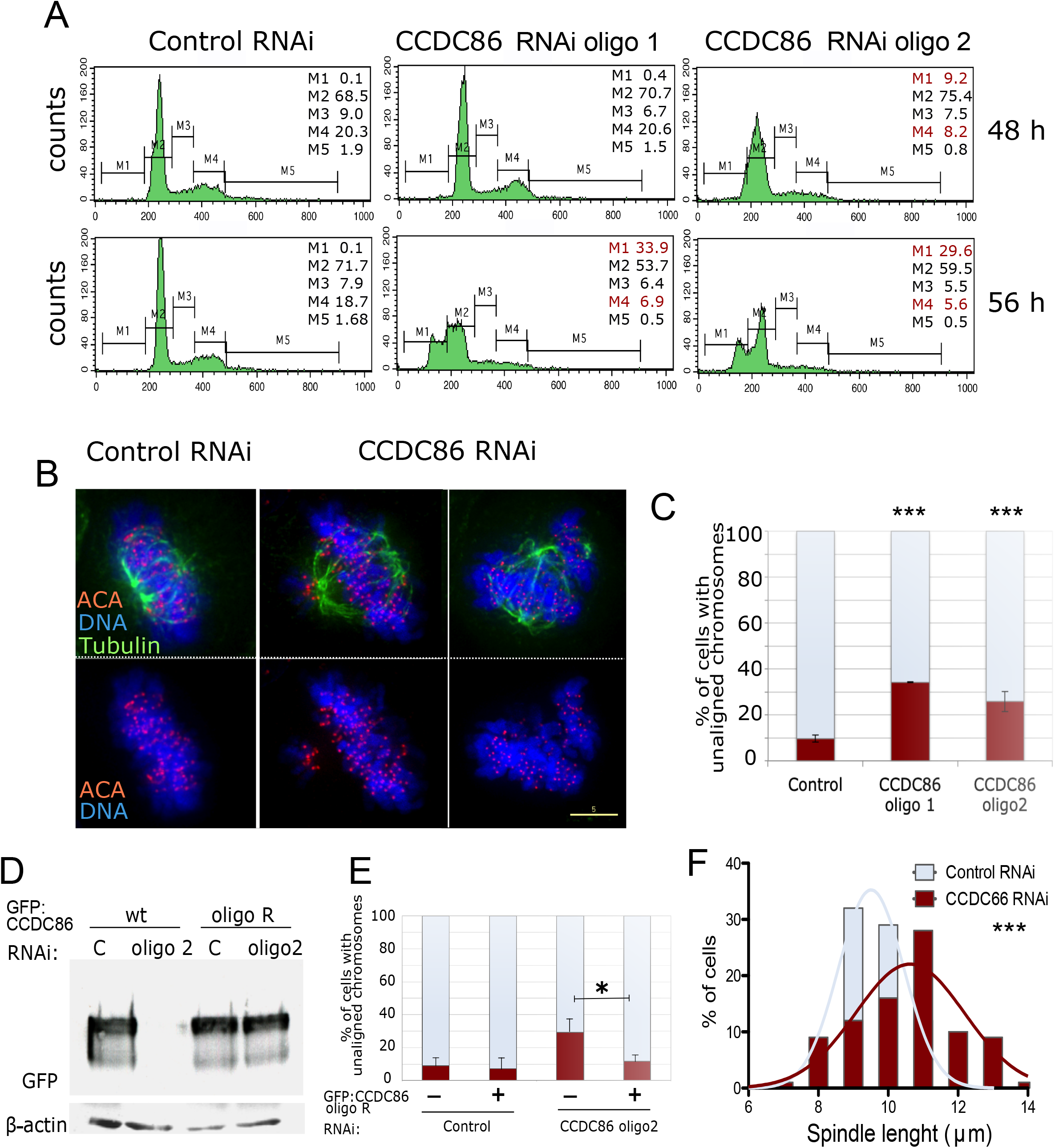
Depletion of CCDC86/Cyclon compromises normal mitotic progression. A) HeLa cells were transfected with Control or CCDC86 siRNA oligos for 48 or 56h. Cells were then harvested and subjected to cell-cycle analyses by FACS. The percentages of cells in each gated interval are shown; M1 (sub-G1), M2 (G1), M3 (S), M4 (G2) and M5 (> 2N). In red are highlighted the stages where there is a significant deviation from the control distribution. B) HeLa cells were treated with either siRNA control or CCDC86 oligos (Oligo #1 or Oligo #2) for 48h and then fixed and stained for tubulin (green), anti-centromere antibody (ACA) (red) and counterstained with DAPI (blue). Representative images of normal aligned metaphase cells (left) and cells with unaligned chromosomes (right) after each RNAi treatment. Scale bar 5μm. C) Quantification of mitotic cells with unaligned chromosomes form the experiment in (B). The graph shows the average or 3 experiments and the error bars represent the SD. N= 166 (Control), 228 (oligo 1) and 182 (oligo 2). Statistical analyses were conducted using the Chi squared test and *** indicates p < 0.0001. D) HeLa cells were transfected with a GFP:CCDC86 wt or the oligo2 resistant construct together with the control or CCDC86 (oligo2) siRNA oligos for 48h. Cells were collected and whole cell lysates were subjected to SDS page and immunoblotting using an anti:GFP or beta actin antibody. E) HeLa cells were treated as in (D) then fixed and immunostained as in (B). The graphs show the quantification of mitotic cells with unaligned chromosomes in the different conditions. The graph shows the average or 3 experiments and the error bars represent the SD. N= 166 (Control untransfected), 130 (Control transfected), 510 (CCDC86 RNAi untransfected), 170 (CCDC86 RNAi transfected). Statistical analyses were conducted using the Chi squared test and * indicates p < 0.05 F) Distribution of the spindle lengths obtained from the experiment in (B). Pole-to-pole distances were obtained from bi-polar metaphase cells when the spindle poles were on the same focal plane. N=35. Statistical analyses were conducted to compare the spindle lengths between Control and CCDC86 RNAi by Mann Whitney test.

We therefore analysed the chromosomal phenotype of control and CCDC86-depleted mitotic cells. This analysis revealed a significant increase in cells where most of the chromosomes were aligned at the metaphase plate together with a few mis-aligned chromosomes (Figure 4B,C). Importantly, this phenotype was rescued by expressing a CCDC86 cDNA resistant to oligo2 (Figure 4D,E). Chromosome attachment errors are also often associated with spindle length variations (Nannas et al., 2014); we therefore measured the spindle length in metaphase cells with a bi-polar spindle. Indeed, CCDC86 RNAi cells have a significant longer spindle length than cells treated with control RNAi (Figure 4F).

All these data together show that CCDC86 contributes towards the execution of an error-free mitosis.

### CCDC86/Cyclon is a MYCN regulated gene with a prognostic value for Neuroblastoma patients

Although very little is known about CCDC86 biology, this protein has already emerged as an important predictor of cancer resistance and outcome in several studies (Bouroumeau et al., 2021) (Emadali et al., 2013). Furthermore, CCDC86 was identified to be an autonomous tumour growth driver that cooperates with MYC to drive aggressive lymphoma growth *in vivo* (Emadali et al., 2013).

Neuroblastoma is a tumour where MYCN amplification correlates with poor prognosis. We therefore analysed survival data for Neuroblastoma patients in the Kokac and ag44kcwolf cohorts using the R2 genomics platform and related to CCDC86 expression. As the Kaplan Meier curves show, high levels of CCDC86 sttrongly correlate with a very low survival rate in this type of cancer (Figure 5 A). We then stratified the patients in each cohort based on the tumours that have MYCN amplified or not and their stage. The results obtained from both cohorts are very similar and clearly show that high expression of CCDC86 is found in patients with MYC amplifications (Figure 5 B, D) and the level of CCDC86 positively correlates with stage progression and malignancy (Figure 5 C and D). In fact, in stage 4s tumours (these tumours are not metastatic) CCDC86 presents low expression. These data suggest that CCDC86 expression is driven by MYCN and that, even in neuroblastoma patients, represent a very useful prognostic marker.

**Figure 5.**
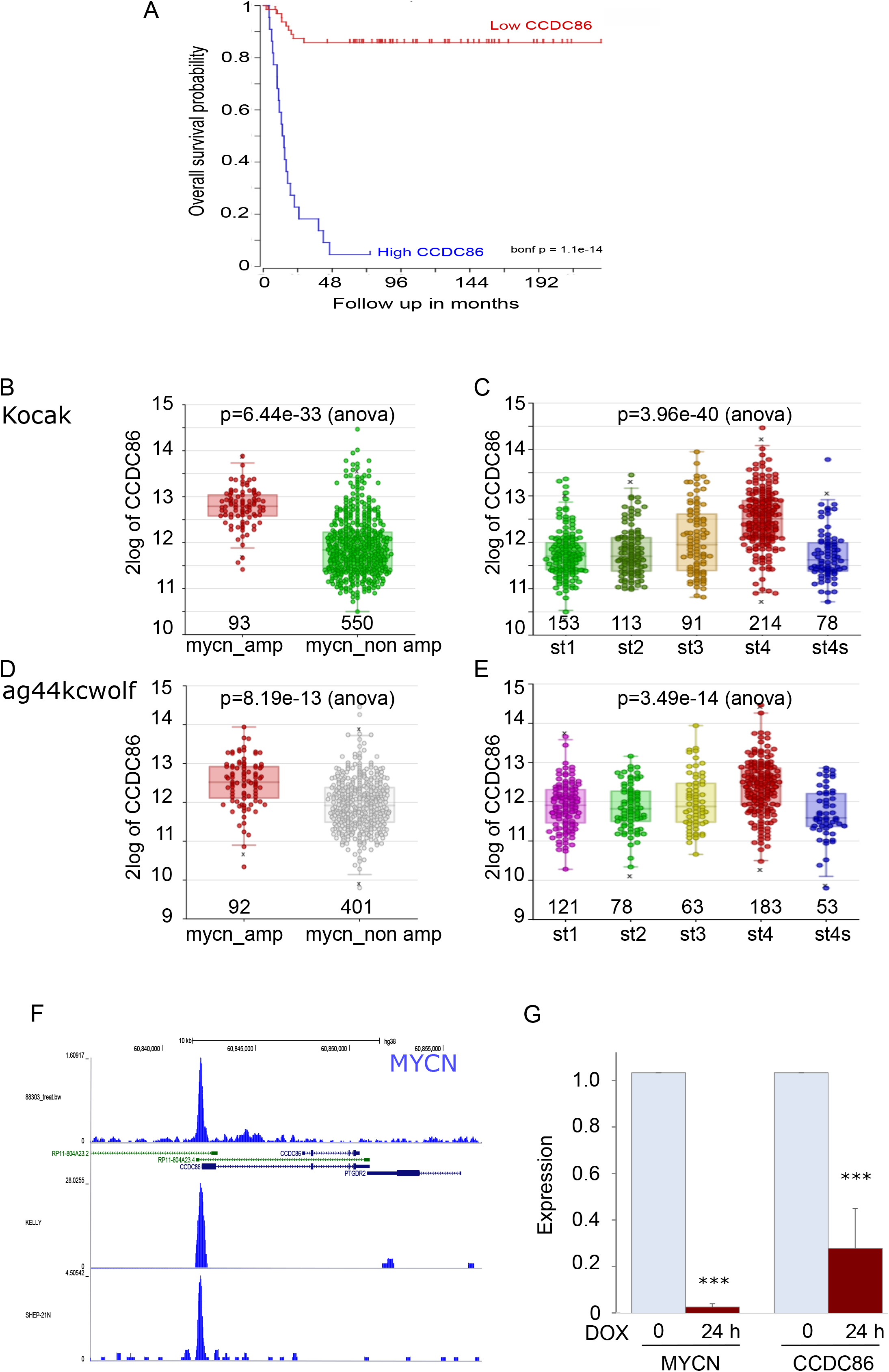
CCDC86 expression is regulated by MYCN represents a prognostic marker for Neuroblastoma patients. A) Kaplan Maier curve (survival probability in months) of neuroblastoma patients expressing low (red curve) or high (blue curve) levels of CCDC86. B) CCDC86 expression levels in neuroblastoma patients with (red) and without (green) MYCN amplification (Kocak cohort). C) CCDC86 expression levels in neuroblastoma patients with different stage cancer (Kocak cohort). D) CCDC86 expression levels in neuroblastoma patients with (red) and without (grey) MYCN amplification (ag44kcwolf cohort). E) CCDC86 expression levels in neuroblastoma patients with different stage cancer (ag44kcwol cohort). F) MYCN ChIP seq profiles at the CCDC86 locus in different neuroblastoma cell lines. G) qPCR analyses of MYCN and CCdc86 expression in TET21-N cells without (0) (light blue bars0 and with 24 h doxycycline (DOX) treatment (brown bars). This represents the average of 3 independent experiments. Error bars indicate the standard deviation. Datasets were analyses by student T test. ***=p<0.0001

We therefore analysed MYCN occupancy at CCDC86 promoters in different MYCN amplified cell lines using the UCSC genome browser. In all profiles analysed, a clear peak for MYCN is present at CCDC86 promoter (Figure 5 F). To confirm the dependency of CCDC86 expression on MYCN we then used the neuroblastoma cell line TET21-N where MYCN expression can be modulated by doxycycline (Lutz et al., 1996). Addition of doxycycline to the culture decreases the expression of MYCN in 24h (Figure 5 G – left) as evaluated by qPCR. Cell viability is not affected by MYC expression in this system. We then tested the expression of CCDC86 in the same conditions; strikingly, repression of MYCN significantly decreases CCDC86 expression. These data therefore confirm that CCDC86 expression is MYCN regulated in neuroblastoma, and it is a useful prognostic marker in this type of cancer.

## Discussion

The chromosome periphery is an outer layer coating the surface of mitotic chromosomes that contains a large number of proteins with diverse functions during interphase. The list of components for this compartment is still growing, as more proteins associated with the chromosome periphery are been identified by proteomics screens of isolated chromosomes (Ohta et al., 2011) (Takata et al., 2007). Despite the ever-growing list of components, the chromosome periphery is not very well characterized either in its composition or in its function. It is also unclear whether the chromosome periphery is functionally a single domain or if it represents an assembly of multiple partially overlapping subcomplexes that collaborate to execute various functions of this structure. The diversity of chromosome periphery associated proteins (Van Hooser et al., 2005) reveals a need for a better understanding of the composition and function of this complex compartment.

We have re-analysis the mitotic chromosome proteome with the goal to identifying novel components of the chromosome periphery (Ohta et al., 2011). This approach highlighted a new chromosomal protein: the Coiled-Coil Domain Containing 86 (CCDC86/cyclon).

CCDC86 was previously described as a downstream effector of IL-3 signaling upon cytokine induction in hematopoietic stem cells (Hoshino and Fujii, 2007). CCDC86 was also shown to play a role in T-cell activation-induced cell death (AICD) (Saint Fleur et al., 2009) and studies have also linked CCDC86 expression with cancer progression (Emadali et al., 2013) (Bouroumeau et al., 2021).

However, CCDC86 biology still remains largely unexplored.

We discovered that CCDC86 exhibits a localisation pattern that in many ways resembles Ki-67 and other chromosomal periphery proteins (Amin et al., 2007) (Ma et al., 2007). A GFP-fusion of CCDC86 localizes to the nucleolus during interphase and is recruited to the chromosome periphery during mitosis. However, CCDC86 exhibits its most prominent association with the chromosome periphery late in mitosis. The weaker association of CCDC86 with the mitotic chromosomes in early mitosis could be the result of a dynamic behavior of the protein when phosphorylated. In fact, CCDC86 presents several phosphorylation sites that are affected by nocodazole treatment (as shown in the phosphosite database https://www.phosphosite.org/homeAction.action).

Ki-67, a major determinant for assembly of chromosome periphery (Booth et al., 2014), controls the recruitment of CCDC86 to the chromosomes and RNAi mediated depletion of Ki-67 prevented CCDC86 association with the chromosome periphery. Interestingly, even though Ki-67 is necessary for CCDC86 recruitment to the chromosomes, it is not sufficient. A mutant version of GFP:CCDC86Δ_63,_ lacking the first AT-Hook domain, abolished the localization of the protein to mitotic chromosomes. This observation could either indicate that CCDC86 has a chromosome periphery targeting module that does not solely depend on a putative Ki-67 interaction or that the N terminus of the protein contains the interaction domain for Ki-67 (or other Ki-67 recruited protein) binding. Combining these results together with the protein gene ontology data and the CCDC86 Protein-Protein Interactome from GPS-Prot, we conclude that CCDC86 is a *bona fide* chromosome periphery associated protein.

CCDC86 depletion led to a novel phenotype in which Ki-67 and Nucleolin were still able to accumulate at the chromosome periphery but also formed abnormal cytoplasmic NDF-like foci. This is the first time Ki-67 has been observed in abnormal NDFs. For Nucleolin, these foci were more abundant and increased in number, compared to control cells and they were atypically present in prometaphase (Ochs et al., 1983).. Together, these observations reveal that upon depletion of CCDC86 the chromosome periphery compartment is maintained, but some of its components exhibit additional abnormal interactions (Figure 6 B).

**Figure 6.**
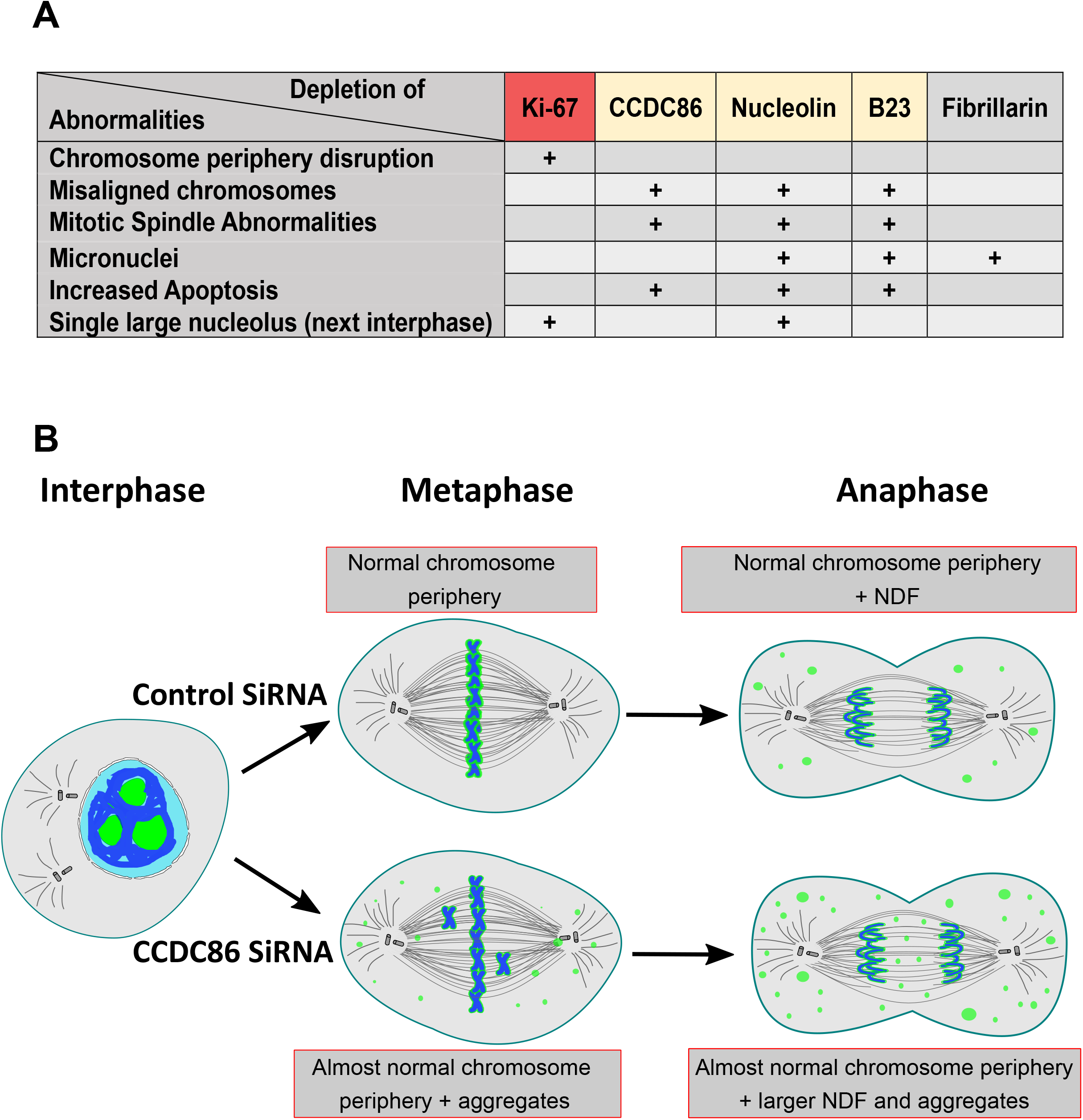
Model for mitotic functions of subcomplexes of the chromosome periphery. A) Table showing the major phenotypes associated with the depletion of the indicated proteins (Amin et al., 2007) (Booth et al., 2014) (Ma et al., 2007) (Ugrinova et al., 2007). B) Graphic summary of CCDC86/Cyclon function in mitosis

Ki-67 association with the chromosome periphery is more dynamic in early mitosis and more stable from mitotic exit onwards (Endl and Gerdes, 2000) (MacCallum and Hall, 2000) (Schluter et al., 1993) (Takagi et al., 2014). During mitosis, Ki-67 is hyperphosphorylated in a CDK1 dependent manner. Those phosphorylations decrease the affinity of Ki-67 for the DNA until anaphase onset, at which point PP1 reverses the phosphorylations, thus increasing Ki-67 association with the chromatin (Saiwaki et al., 2005) (Takagi et al., 2014) and may play a role inregulating Ki-67 surfactant properties (Cuylen et al., 2016) (Stamatiou and Vagnarelli, 2021). Staurosporine treatment of metaphase arrested cells causes disassociation of Ki-67 and B23 from the chromosome periphery and formation of abnormal cytoplasmic foci (NDF-like foci) (MacCallum and Hall, 2000). Interestingly, CCDC86 depletion leads to a similar phenotype. We hypothesize that Ki-67 dynamics may not only be regulated by its phosphorylation status but also by specific protein-protein interactions.

We observed errors in chromosome alignment in cells depleted of CCDC86. This phenotype was specific as it was rescued by transient expression of siRNA-resistant CCDC86. In addition, cells lacking CCDC86 displayed a significant longer spindle length compared to control cells and an increase in apoptosis, possibly a consequence of mitotic defects. These observations, combined with the partial disassociation of Nucleolin from chromosomes after CCDC86 depletion, suggest that chromosome misalignment and mitotic spindle defects observed in these cells may be consequences of the dissociation of Nucleolin and, to a certain extent, of B23 from chromosomes. In fact, depletion of Nucleolin or B23 lead to chromosome misalignment, defects in mitotic spindle formation and apoptosis (Amin et al., 2007) (Ma et al., 2007) (Ugrinova et al., 2007); Nucleolin appears to be upstream of B23, as B23 disappears from the chromosome periphery after nucleolin depletion (Ma et al., 2007). Our data are consistent with CCDC86 being upstream of B23 for the regulation of mitotic spindle formation and correct microtubule attachment.

Interestingly, these phenotypes are masked when Ki-67 is depleted, and the entire chromosome periphery layer is removed. Being able to remove only a subset of proteins can thus reveal some different functions that are embedded in this compartment (Figure 6A). The classifications of the phenotypes seem to suggest the existence of a cluster of functions for the chromosome periphery subcomplexes. In fact, the recently identified NWC (NOL11-WDR43-Cirhin) complex, which is part of the mitotic chromosome periphery is required for the centromeric enrichment of Aurora B and the downstream phosphorylation of histone H3 at threonine 3 (Fujimura et al., 2020).

We have also shown that CCDC86 is a MYCN driven protein that with prognostic value in Neuroblastoma. In this respect, considering that CCDC86 depletion induces apoptosis, it would be interesting to evaluate in the future if this protein could also represent a clinically relevant drug target.

Our study represents a further step towards understanding the complexity and functional significance of the chromosome periphery in mitosis and cell-cycle progression.

## Competing interests

All the Authors declare No competing interests.

## Funding

The Vagnarelli lab is supported by the Wellcome Trust Investigator award 210742/Z/18/Z to Paola Vagnarelli. The study on MYC (Figure 5) was supported by Kidscan PhD studentship awarded to PV.

The Ohta lab is supported by Research Foundation for the Electrotechnology of Chubu (grant number R-01227) to SO.

Work in the Earnshaw laboratory is funded by Wellcome, from whom WCE holds a Principal Research Fellowship (grant no. 107022).

## Data availability

The datasets used for this study are:

1. For the mitotic chromosome analyses: (Ohta et al., 2010)
2. For the neuroblastoma patients, data sets are available at https://hgserver1.amc.nl/cgi-bin/r2

Imaging and quantification datasets are available upon request from the corresponding authors and will be shared via Figshare.

## Notes

### Competing Interest Statement

The authors have declared no competing interest.

